# The Influence of Material Surface Properties on Initial Bacterial Attachment and Biofilm Formation on Built Environment Surfaces

**DOI:** 10.64898/2026.01.28.702373

**Authors:** Kobi Talma, Nathan Bossa, Evan Hankinson, Lijia Gao, Aicha El Kharraf, Mark R. Wiesner

## Abstract

Biofilms in the built environment (BE) can harbor pathogens and have been linked with negative health outcomes, particularly in hospital environments. The formation of biofilms requires bacterial cell attachment on surfaces, such as hospital plumbing, which can have varying properties, including roughness, wettability, chemistry, and charge. Despite the importance of bacterial attachment to surfaces, the role of multiple surface properties has been minimally investigated. Using seven materials with differing surface characteristics, this work considers the initial attachment of *Escherichia coli*, *Pseudomonas aeruginosa*, *Bacillus subtilis,* and *Staphylococcus aureus* to investigate the impact of several surface characteristics. Biofilm formation was evaluated to determine the impact of surface characteristics on longer-term attachment. The initial attachment of all bacterial species was not significantly influenced by material surface properties, with similar attachment seen across materials tested. Bacterial cell envelope morphology affected attachment, with Gram-negative species displaying greater attachment than Gram-positive species. Initial attachment was not related to biofilm formation, which was highest on stainless steel surfaces while differences between bacterial species remained.

**SYNOPSIS:** Limiting pathogen presence in the Built Environment is key to limiting risk to public health. This study reports that bacterial properties, not material properties, have greater impact on attachment to surfaces, indicating the importance of a microbial-focused effort to engineer the microbiome of the Built Environment.

**TABLE OF CONTENTS (TOC) GRAPHIC:** 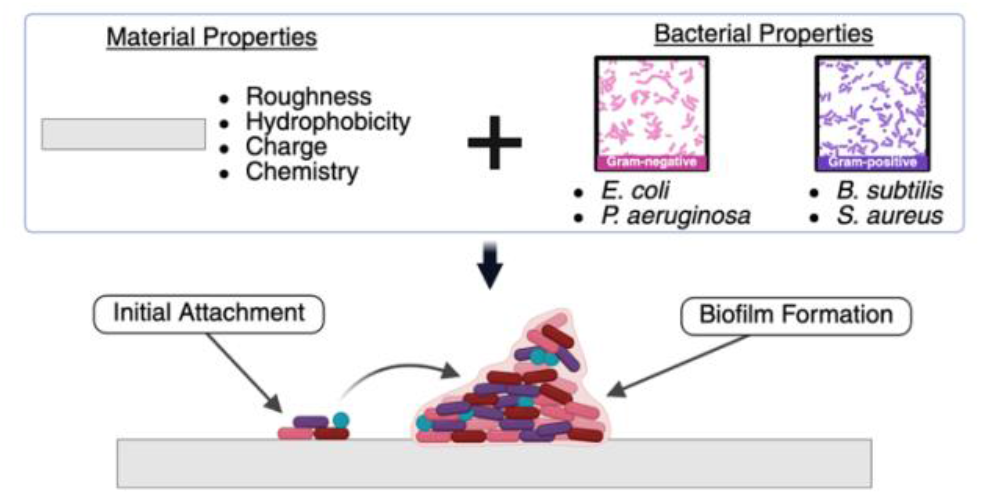

## INTRODUCTION

Pathogen presence in the built environment (BE) poses a risk to human health^1^. This is particularly relevant in settings such as hospitals, where hospital-acquired infections represent a significant burden on the healthcare system. The estimated annual cost of hospital-acquired infections is between $7.2 – 14.9 billion^2,3,4^ in the United States healthcare system and results in significant morbidity and mortality within the United States^5,6^. Premise plumbing has been shown to support the growth of pathogenic bacteria^7,8^ and contribute to between 7% and 40% of infections^9,10^. This pathogen growth often occurs in the form of biofilms, which can serve as a protective barrier in extreme environments^11^. For example, bacteria established in hospital plumbing may migrate from these biofilm reservoirs up towards the sink drain, where droplet dispersion during faucet operation can lead to infection^12^. The formation of bacterial biofilm involves several steps: (1) attachment, (2) microcolony formation, (3) matrix formation, (4) macrocolony formation, also known as maturation, and (5) dispersal^13^. The attachment of bacterial cells to surfaces is therefore an integral step to the establishment and reestablishment of biofilms. The use of antimicrobial chemicals, including those to coat surfaces, may increase the prevalence of antimicrobial resistance^14,15^. The current work explores the impacts of surface properties on bacterial attachment with the possibility of manipulating surface properties to limit bacterial attachment as a chemical-free strategy for combating biofilm formation without increasing antimicrobial resistance.

Bacterial attachment and biofilm formation have been studied previously in various contexts, including medical devices^16,17,18^, membrane biofouling^19^, and marine biofouling^20^. Surface properties play a role in attachment through attractive (van der Waals) and repulsive (electrostatic) forces. Although numerous studies have investigated the impact of material properties on bacterial attachment or adhesion, most of these studies have focused on the impact of a single surface property, and efforts to assess the impact of multiple surface properties are essential to advance knowledge^21^. Environmental sampling of the BE has largely failed to consider the impact of any material property on the microbes of the BE, let alone multiple properties^22^. This further amplifies the need to understand the collective impact of multiple surface properties on bacterial attachment and the microbiome of the BE.

In previous studies, the quantification of bacterial adhesion or attachment has been studied using batch tests, where bacteria were put in contact with materials for a defined time period, and cells remaining on the materials’ surface after rinsing were quantified. While such an approach yields useful information concerning the tendency of bacteria to establish themselves on a given surface, it does not provide kinetic information on the attachment process and is far from real-life conditions (i.e., fluid dynamics, high contact surfaces, limited nutrients). Alternatively, the concept of attachment efficiency has been used extensively in colloid science to quantify the affinity of particles for surfaces in a kinetic context, where α can have values between 0 and 1, and describes the likelihood of a particle attaching upon interacting with a surface. This approach has been used for predicting particle removal in water treatment^23,24,25^, bacterial transport in bioremediation^26^, and the fate and transport of nanomaterials^27,28,29,30^. Surface attachment efficiency, α, is the ratio between the number of successful attachments and the number of collisions with the surface and is typically a function of surface properties (of both the particle and the surface) and environmental conditions. Combined physical and hydrodynamic models for particle motion, α, can be used to determine rates of attachment in porous media^31^. It has also been used to determine aggregation rates^32^, with applications in natural^33,34^ and engineered systems^35^.

In this study, we explore the initial attachment of bacteria to relevant, well-characterized BE materials to fill the gap in assessing multiple surface characteristics’ impact on bacterial attachment. We used model bacteria (*Escherichia coli* K-12 and *Bacillus subtilis* ATCC1174) and environmentally isolated bacteria (*Pseudomonas aeruginosa* and *Staphylococcus aureus*) to i) investigate the effect of material properties on attachment efficiency (α); ii) investigate the effect of bacterial properties/cell envelope morphology on initial attachment efficiency (α); and iii) investigate the effect of material on longer-term biofilm formation.

## MATERIAL AND METHODS

### Material Preparation

Pre-production pellets of six polymers: acrylonitrile butadiene styrene (ABS), high-density polyethylene (HDPE), high-impact polystyrene (HIPS), polycarbonate (PC), polyvinyl chloride (PVC), and polyvinylidene fluoride (PVDF); and stainless steel (SS) spheres were used in this work. ABS, HDPE, HIPS, PC, and PVDF were purchased from McMaster Carr (Elmhurst, Illinois). PVC was purchased from Alphagary (Pineville, North Carolina). SS was purchased from MSE Supplies (Tucson, Arizona). To prepare the materials, 250mL batches of pellets/spheres were washed in 70% ethanol. After ethanol washing, the pellets/spheres were rinsed thoroughly with deionized (DI) water three times. Pellets/spheres were then air-dried in a laminar flow hood and stored in an autoclaved, airtight container until use.

### Material Characterization

The water contact angle (WCA) was measured to determine the surface wettability of the materials, using the sessile drop method with an optical tensiometer (Attension Theta Flex, Biolin Scientific; Phoenix, Arizona). In short, a 4µL drop of DI water was placed on the sample pellet, and images of the drop were captured for 10 seconds, at 33 frames per second, for analysis by the Biolin Scientific software. The average contact angle from all 330 images collected was reported as the contact angle for one replicate. Eight technical replicates were performed for each material type.

Surface roughness was measured by 3D optical/ laser confocal profilometry (VK-X3000, Keyence; Itasca, Illinois) using a 50x objective, with images 208µm × 278µm collected for each material type. For cylindrical pellets (ABS, HIPS, and PC), images were taken of both the flat surface (top of cylinder) and the curved surface (lateral surface). The roughness of cylindrical pellets (ABS, HIPS, and PC) is reported as the weighted average of the flat and curved surfaces of the materials, assuming the flat surface covers one-third of the total surface area, and the curved surface covers the other two-thirds of the total surface area.

The surface charge was measured as zeta potential using a solid surface zeta potentiometer (SurPass, Anton Paar; Austria). The surface charge was measured across a pH range from 2.7 to 7.8, a Boltzmann curve was fit to the zeta potential data, and the zeta potential at pH 7.5 was determined. A Fourier transform infrared (FTIR) spectrophotometer (Nicolet iS50, ThermoFisher Scientific; Waltham, Massachusetts) with an attenuated total-reflection diamond crystal accessory (ATR) was used to evaluate surface functional groups. Thirty-two scans were performed over the spectral range of 3600-400 cm^-1^, and spectra processing was performed using Spectragryph^36^ to identify surface functional groups.

### Bacterial Preparation and Characterization

The four model organisms used in this study were *Escherichia coli* K-12 (*E. coli*), an environmental isolate of *Pseudomonas aeruginosa* (*P. aeruginosa*) from a hospital sink P-trap, *Bacillus subtilis* ATCC 1174 (*B. subtilis*), and an environmental isolate of *Staphylococcus aureus* (*S. aureus*) from a hospital sink basin. The model organisms were grown in liquid cultures in Lennox Luria Broth (LB) media (Sigma-Aldrich; St. Louis, Missouri) to a stationary phase at 37°C with shaking. After reaching the stationary phase, the liquid cultures were centrifuged in 50mL tubes at 3000xg and 4°C for 10 minutes. The nutrient media was aspirated from the tubes, and the pelleted bacteria were resuspended in EPA Moderately Hard water (EPA Mod Hard).

Cell surface charge was measured by zeta potential, calculated from electrophoretic mobility measurements using a Malvern Zetasizer Nano ZS (Malvern; United Kingdom). Cell suspensions were prepared as described above, and measurements were performed in EPA Mod Hard (approximately 2 mM). Cell hydrophobicity was determined using water contact angle measurements using the sessile drop method on bacterial lawns^37,38^ with an optical tensiometer (Attension Theta Flex, Biolin Scientific; Phoenix, Arizona). Bacteria cells were prepared as described above and bacterial lawns were created by filtering bacterial suspensions through sterile mixed cellulose esters (MCE) membrane (Simsii^™^; Issaquah, Washington) of 0.22µm pore size until the fluid had completely passed through the filter. Filters with bacterial lawns were air-dried under sterile conditions for 30 minutes before contact angle measurements were performed.

### Bacterial Attachment in Dynamic Column Conditions

The column testing setup was modified from Rogers et al.^31^ and was organized as follows: dry pellets/spheres were packed into a glass column (25mm x 150mm, Diba Omnifit EZ Chromatography Column, Cole Parmer; Vernon Hills, Illinois) to the 11mm line. Deionized water was pumped into the column from the bottom by a syringe pump until the column was filled with water. The column effluent passed into an in-line ultraviolet-visible spectrophotometer (Evolution 201 UV-visible Spectrophotometer, CAT# 912A0883, ThermoFisher Scientific; Waltham, Massachusetts). The bacteria sample, with a starting optical density at 600nm (OD600) of 0.6, was passed directly into the spectrophotometer, bypassing the column by means of T-connectors, to obtain a maximum absorbance at 600nm for the sample (the initial bacteria concentration, C_0_). The tubing was flushed with deionized water to remove all of the bacteria from the system. Then, roughly 3 pore volumes of the background solution (EPA Mod Hard) were passed through the column to equilibrate the packing material with the background. The bacteria sample was passed through the column at a flow rate of 2.95mL/min, and the absorbance was recorded at approximately every 10 seconds for at least 4 pore volumes.

A tracer study using potassium nitrate was performed to confirm the integrity of the column. Column experiments were also carried out using positively charged aminated 1µm silica particles (nanoComposix; San Diego, California) to calculate α. The details of the tracer study and positive particle column experiments are included in the Supporting Information.

### Calculation of Attachment Efficiency (α)

Colloid filtration theory was used to describe bacterial attachment using α. α can be expressed as a function of experimentally observed concentrations of particles in the column effluent (C) and influent (C_0_)^39^:

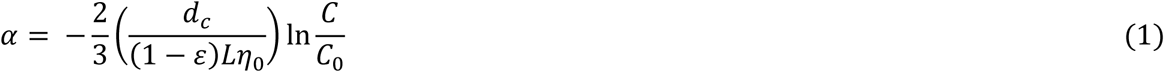

where *d_c_* is the diameter of the collector (pellet/sphere), ε is the porosity of the column, *L* is the column length, and *η*_0_ is the predicted single-collector contact efficiency (calculation described in Supporting Information).

Using the aminated 1µm silica particles as an assumed α = 1 particle, due to their positive charge, the value of α was determined using Eq. 2, with the values of C/C_0_ taken from the steady-state phase of the column data. Experiments for α were conducted in duplicate, and the standard error is reported to compare between experiments.

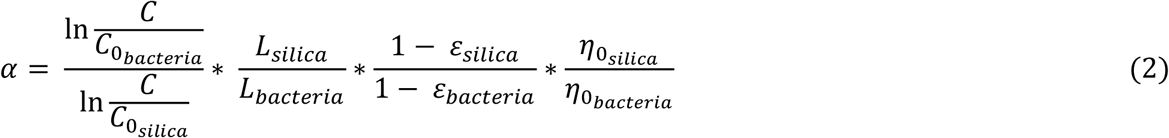

### Evaluation of Biofilm Formation

Crystal violet biofilm assays were done to evaluate the extent of biofilm formation on the material tested. A diluted bacterial solution was prepared in 1/2x R2A media (HiMedia; Kennet Square, Pennsylvania) by adding 20µL of the bacterial solution, starting OD600 = 0.6, to 20mL of diluted minimal nutrient media (1/2x R2A). A known mass of the test material was incubated with 500µL of diluted bacterial solution at 25°C in a 48-well Falcon® plate (Corning; Glendale, Arizona). After incubation for 24 hours, planktonic cells were removed, and the material was rinsed twice with PBS to remove loosely attached cells. Then, the material was stained with 0.1% crystal violet for 15 minutes at room temperature. The crystal violet was removed, and the material was rinsed until no excess crystal violet remained. The material was allowed to dry overnight before the crystal violet was solubilized with 30% acetic acid. The solubilized crystal violet was transferred to a new microplate, and the absorbance was measured at 585nm^40^ using a microplate reader (SPECTROstar Nano, BMG Labtech; Germany).

The measured crystal violet absorbance was normalized using the material surface area using the following formula:

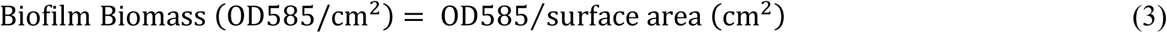

### Statistical Analysis

Statistical analysis considered the α results and biofilm results as a function of material, organism, and the interaction between material and organism. A two-way analysis of variance (ANOVA) was performed in R to test for significance. Where results were statistically significant, post hoc pairwise comparisons were performed using the Bonferroni correction. A p-value of <0.05 was considered to be statistically significant. The tabulated results of the statistical tests are included in the Supporting Information.

## RESULTS AND DISCUSSION

### Characterization of Material Surface Properties

The materials were selected for their range of expected properties and prevalence in the BE. HDPE and PVC are commonly used for pipe materials in both distribution systems and premise plumbing^41^. PVDF is used in high-purity water systems, such as Ultrapure water, due to its chemical and biological stability^42^. ABS is used in drain-waste-vent systems, which include the drainpipes from sinks and other plumbing in the BE^43^. HIPS is found in many plastic surfaces in the BE, particularly in packaging applications^44^. PC is another common plastic material in the BE, with more resistance than HIPS^45^. SS was selected as a control material and is often used due to its corrosion-resistant properties^44^.

The materials were characterized for several surface properties to identify any differences that may influence bacterial attachment, and as shown in Table 1, the materials studied presented different surface properties. The properties that have been the focus of previous studies investigating bacterial attachment to surfaces are water contact angle^46,47^, roughness^47,48^, and surface charge^17,49^.

**Table 1:**
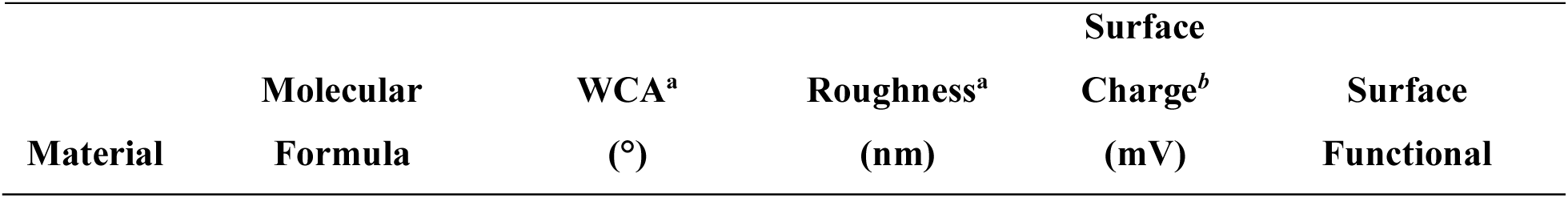
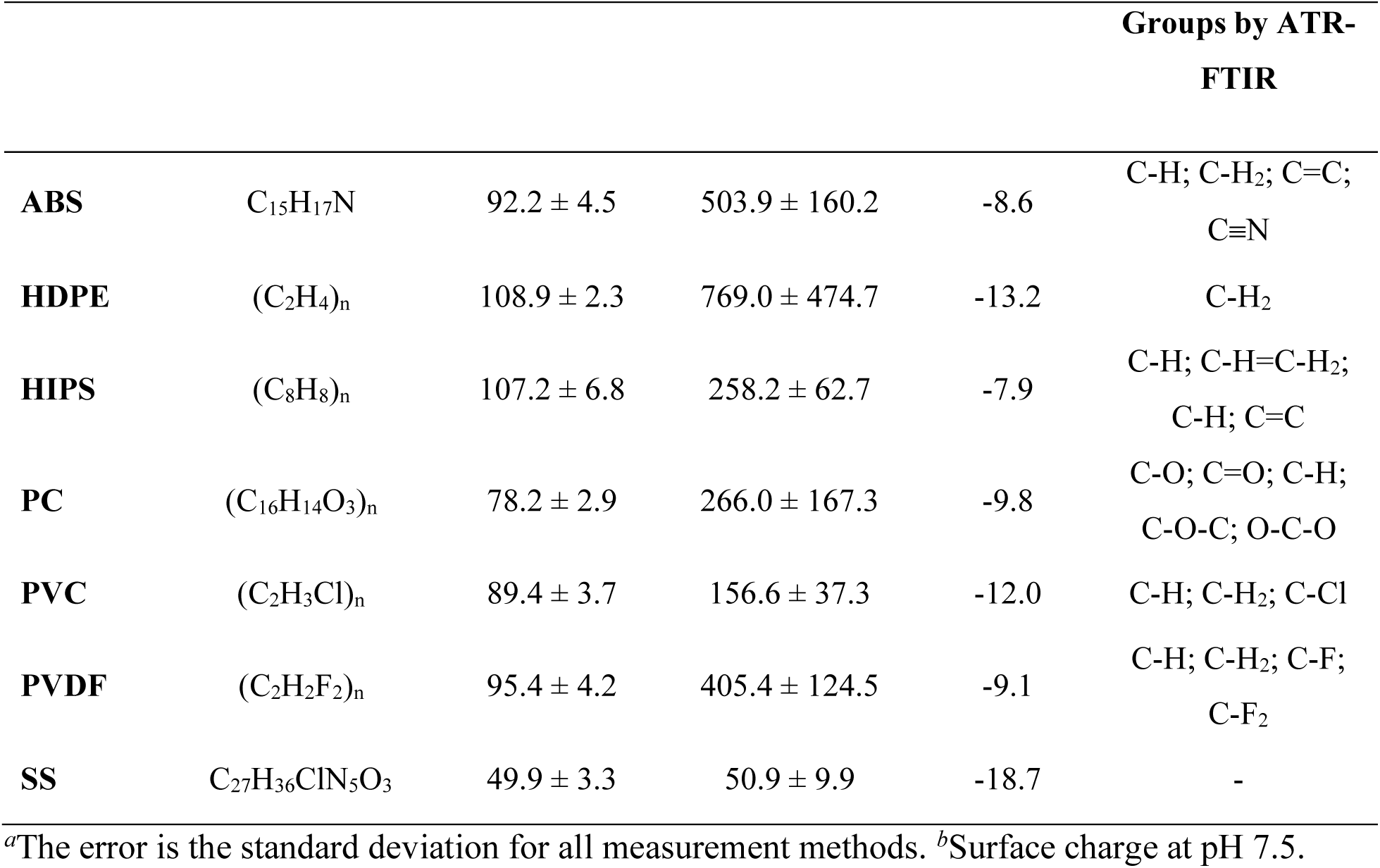
Material Properties and Surface Characteristics.

Surfaces with water contact angles greater than 90° are generally considered hydrophobic, while those with water contact angles less than 90° are operationally defined as hydrophilic^50^. The selected materials range from hydrophilic (SS) to hydrophobic (HDPE and HIPS), with several materials falling near the 90° threshold. Surface roughness ranged from very smooth (SS = 50.9 ± 9.9 nm) to rough (HDPE = 769 ± 474.7 nm). The standard deviation for the roughness measurements describes the heterogeneity of the surfaces, with regions of particularly high roughness visible on some surfaces (Figures S.5.1-S.5.2). The surface charge was negative for all tested materials, ranging between –7.9 mV and -18.7 mV. The surface chemistry was measured using ATR-FTIR, and the ATR-FTIR spectra are provided in Figure S.5.3 in the Supporting Information. The plastics evaluated in this study (ABS, HDPE, HIPS, PC, PVC, and PVDF) exhibited a range of functional groups, consistent with their polymer composition, which is shown in Table 1. ABS, HDPE, and HIPS surfaces showed non-polar groups, with ABS also showing nitrile (C≡N) groups associated with acrylonitrile. PC presents more oxygen-containing surface functionality that makes the surface more polar, which corresponds with the lower water contact angle than the other plastics. PVC and PVDF surfaces are chemically distinct, with the presence of chlorine and fluorine on the surface, respectively. Stainless steel lacks organic functional groups, explaining the lack of signal found in the ATR-FTIR measurement of SS. Interestingly, the material properties measured in this study showed that while all materials exhibited negative zeta potentials, the most hydrophilic material (SS) had the lowest surface roughness, while the material with the greatest roughness (HDPE) was the most hydrophobic.

### Effect of Material Surfaces on Initial Bacterial Attachment

The parameter α was used to evaluate the initial attachment of bacteria to the selected BE materials. Two lab-derived bacteria (*E. coli* and *B. subtilis*) and two bacteria isolated from the hospital environment (*P. aeruginosa* and *S. aureus*) were selected for this study to compare the initial attachment behavior of bacteria with different cell morphology (Gram-negative vs. Gram-positive) and different pathogenicity (lab-derived vs. hospital isolates).

To determine the value of α for bacterial cells attaching to BE materials, column experiments were performed for all four bacterial species using the seven different characterized materials. Breakthrough curves, where the bacterial cells become observable in the column effluent, are shown in the Supporting Information. The attachment efficiency to the seven materials was evaluated for all four bacterial species (*E. coli*, *P. aeruginosa*, *B. subtilis*, and *S. aureus*). The measured normalized concentrations (C/C_0_) and calculated α values are shown in Table 2. For all materials and bacteria, α is low with values below 0.05.

**Table 2:** Calculated Values of α for Four Bacterial Species to Various Materials.

| Bacteria | Material | $C/C_0^a$ | $\alpha^{a,b}$ |
| --- | --- | --- | --- |
| *E. coli* | ABS | $0.84 \pm 0.054$ | $0.033 \pm 0.011$ |
| | HDPE | $0.86 \pm 0.046$ | $0.029 \pm 0.012$ |
| | HIPS | $0.83 \pm 0.027$ | $0.033 \pm 0.0079$ |
| | PC | $0.87 \pm 0.039$ | $0.025 \pm 0.0075$ |
| | PVC | $0.86 \pm 0.023$ | $0.025 \pm 0.0042$ |
| | PVDF | $0.88 \pm 0.041$ | $0.034 \pm 0.013$ |
| | SS | $0.91 \pm 0.014$ | $0.020 \pm 0.0034$ |
| *P. aeruginosa* | ABS | $0.84 \pm 0.0074$ | $0.030 \pm 0.0011$ |
| | HDPE | $0.86 \pm 0.0038$ | $0.025 \pm 0.00080$ |
| | HIPS | $0.84 \pm 0.0030$ | $0.028 \pm 0.00012$ |
| | PC | $0.87 \pm 0.0034$ | $0.025 \pm 0.0017$ |
| | PVC | $0.86 \pm 0.040$ | $0.022 \pm 0.0065$ |
| | PVDF | $0.90 \pm 0.0045$ | $0.030 \pm 0.0016$ |
| | SS | $0.87 \pm 0.0071$ | $0.025 \pm 0.0014$ |
| *B. subtilis* | ABS | $0.98 \pm 0.014$ | $0.0036 \pm 0.0031$ |
| | HDPE | $0.97 \pm 0.013$ | $0.0078 \pm 0.0039$ |
| | HIPS | $1.00 \pm 0.0016$ | $0.00031 \pm 0.00032$ |
| | PC | $0.98 \pm 0.017$ | $0.0034 \pm 0.0035$ |
| | PVC | $0.99 \pm 0.00056$ | $0.0025 \pm 0.000064$ |
| | PVDF | $0.94 \pm 0.037$ | $0.021 \pm 0.012$ |
| | SS | $1.00 \pm 0.0033$ | $0.00073 \pm 0.00072$ |
| *S. aureus* | ABS | $0.96 \pm 0.0024$ | $0.0077 \pm 0.00050$ |
| | HDPE | $0.95 \pm 0.0011$ | $0.0086 \pm 0.00040$ |
| | HIPS | $0.97 \pm 0.0051$ | $0.0055 \pm 0.00068$ |
| | PC | $0.93 \pm 0.031$ | $0.014 \pm 0.0058$ |
| | PVC | $0.96 \pm 0.0046$ | $0.0068 \pm 0.00050$ |
| | PVDF | $0.93 \pm 0.018$ | $0.021 \pm 0.0051$ |
| | SS | $0.95 \pm 0.0083$ | $0.010 \pm 0.0018$ |

<sup>a</sup>The error is the standard error based on duplicate measurements. <sup>b</sup> $\alpha$ values were compared for columns of the same bacterial species.

The low attachment found during this study can be explained by repulsive electrostatic forces between negatively charged surfaces and negatively charged bacterial cells^51,52^. Column experiments and attachment efficiency (α) have been used previously to evaluate the transport and attachment of bacteria in porous media^53,54,55^. McCaulou et al. found α values between 0.04 and 0.4 for the interaction between bacteria and different quartz surfaces^53^. The largest α values found were between negatively charged bacteria and coated surfaces with a positive charge. Deshpande et al. showed that *P. fluorescens* had an α value of 0.094 when interacting with silica surfaces^54^. Increased ionic strength increased the attachment of *P. fluorescens*, up to 0.206 at ionic strength of 3 x 10^-2^M, which has also been seen with attachment of other biological particles^31,55^.

Usually, hydrophilic surfaces, with increased wettability, led to increased attachment in previous studies. Bruzaud et al. found that more wild-type *P. aeruginosa* cells attached to a stainless steel surface with WCA of 61° (attached cells ≅ 9 x 10^4^ CFU/cm^2^) than to a superhydrophobic stainless steel with WCA of 147° (attached cells ≅ 3 x 10^3^ CFU/cm^2^)^46^. Similarly, Liu et al. found increased attachment of *E.* coli on hydrophilic titanium (WCA = 42.0°) and stainless steel (WCA = 65.8°) surfaces, compared to hydrophobic Ni-P-PFTE surfaces (WCA 108.0° and 117.8°)^56^. Conversely, another study found that the near-wall velocity of bacteria was reduced when interacting with hydrophobic surfaces, promoting adhesion to the surface^57^. Rough surfaces have also been shown to lead to increased microbial attachment. Mu et al. showed that bacterial densities were low on smooth surfaces, with values between 2-3 x 10^-2^ cells/µm^2^, but as roughness increased, bacterial density increased to 5-10 x 10^-2^ cells/µm^2^ ^47^. Yang et al. found a positive correlation between bacterial attachment and surface roughness above a threshold of 6 nm^58^. However, according to two-way ANOVA, changes in material had no significant effect on α (Table S.6.1). Additionally, when bacterial attachment is plotted against the surface properties of the material, no trend is observed (Figure S.8.1). Overall, the results showed that material properties in this study did not significantly impact initial bacterial attachment, contrary to previous studies.

### Effect of Bacterial Cell Properties on Initial Attachment

The cell surface charge and hydrophobicity of the four bacterial species were measured, and the results are shown in Table 3, along with the cell envelope morphology and pathogenicity.

**Table 3:**
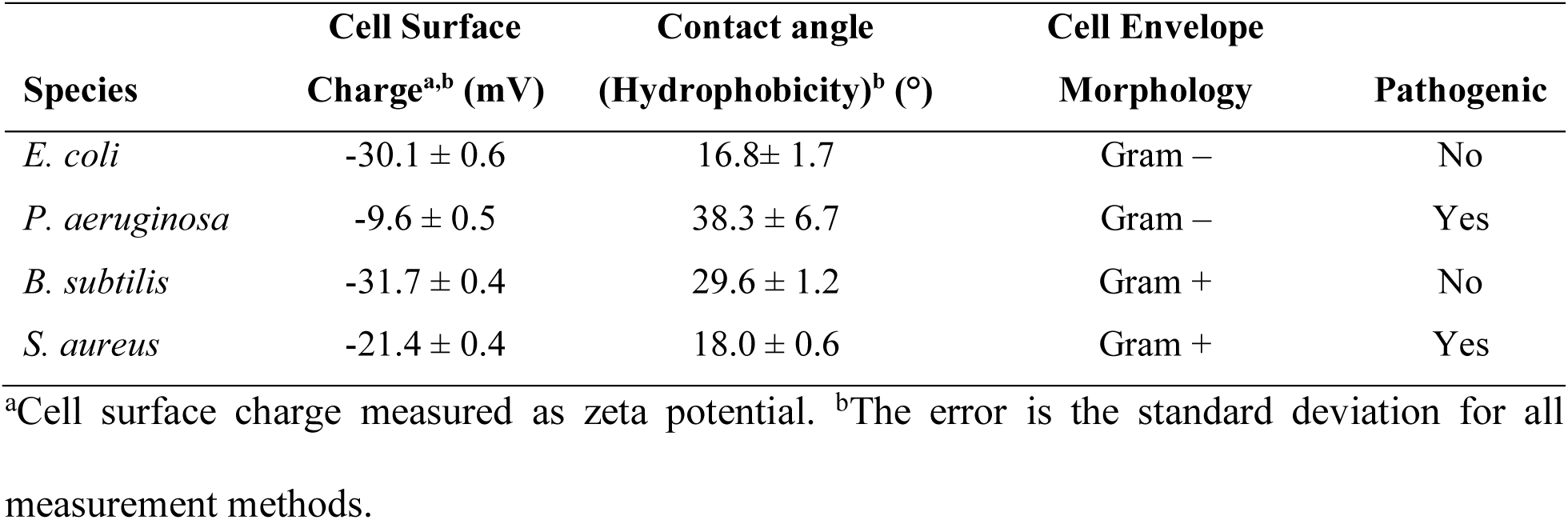
Bacterial Cell Surface Characteristics and Pathogenicity.

| Species | Cell Surface | Contact angle | Cell Envelope |  |
| --- | --- | --- | --- | --- |
|  | Charge<sup>a,b</sup> (mV) | (Hydrophobicity)<sup>b</sup> (°) | Morphology | Pathogenic |
| *E. coli* | -30.1 ± 0.6 | 16.8 ± 1.7 | Gram – | No |
| *P. aeruginosa* | -9.6 ± 0.5 | 38.3 ± 6.7 | Gram – | Yes |
| *B. subtilis* | -31.7 ± 0.4 | 29.6 ± 1.2 | Gram + | No |
| *S. aureus* | -21.4 ± 0.4 | 18.0 ± 0.6 | Gram + | Yes |

<sup>a</sup>Cell surface charge measured as zeta potential. <sup>b</sup>The error is the standard deviation for all measurement methods.

Figure 1 shows the results for α compared between bacterial species when pooled across all seven materials. Two-way ANOVA results showed that species had a significant effect on α (Table S.6.1) and post-hoc pairwise comparisons showed statistically significant differences in α between both Gram-negative species (*E*. *coli* and *P. aeruginosa*) and both Gram-positive species (*B. subtilis* and *S. aureus*) (Table S.6.2). The α results did not show a dependence on any of the other bacterial cell properties (surface charge, hydrophobicity, pathogenicity). The initial attachment efficiency of the Gram-negative species was found to be higher than the initial attachment efficiency of the Gram-positive species by an order of magnitude (0.029 and 0.026 for *E. coli* and *P. aeruginosa* compared to 0.0056 and 0.011 for *B. subtilis* and *S. aureus*).

**Figure 1:**
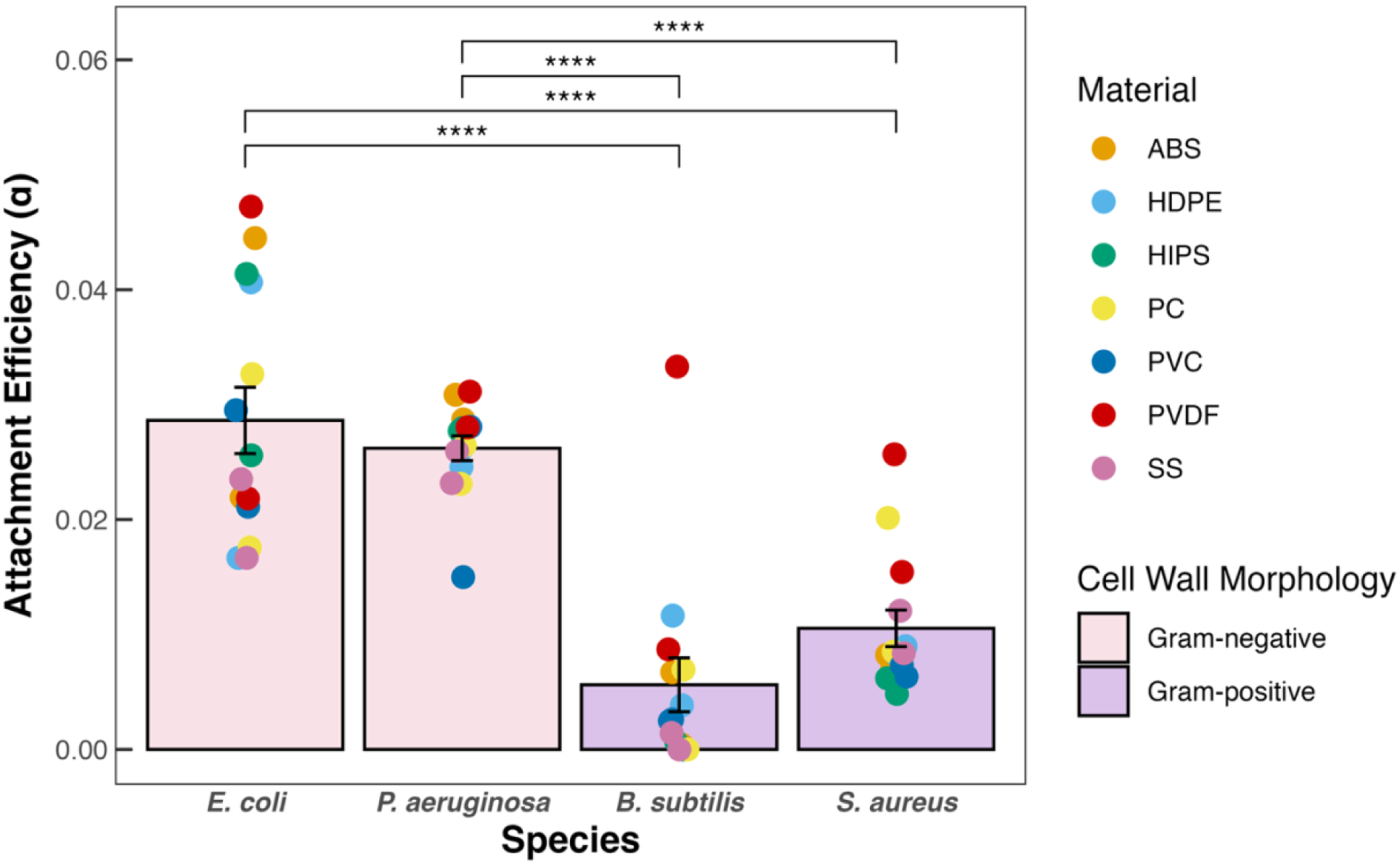
Average bacterial attachment efficiencies for different bacterial species across all materials tested. Error bars represent the standard error from 14 measurements. Asterisks indicate significant differences among groups based on Bonferroni-adjusted pairwise comparisons: *p* ≤ 0.0001 (****).

Gram-positive bacteria have a thick peptidoglycan layer that increases the rigidity of their cell envelope^59^. This rigidity may limit initial attachment due to an inability of the cell envelope to adapt to nanoscale roughness present on the surface, leading to sub-optimal surface contact. The rigid peptidoglycan layer of Gram-positive bacteria limits mechano-selectivity towards a surface, compared to the more sensitive surface structures of Gram-negative bacteria^60^. The effect of cell envelope rigidity was also shown in a study comparing the effect of zero-valent iron, where the envelope of Gram-negative bacteria was more susceptible to the zero-valent iron adhering to the cell membrane than that of Gram-positive bacteria^61^. Additionally, Gram-negative bacteria have external lipopolysaccharides and adhesin proteins on their cell walls, which facilitate attachment to surfaces^62^.

### Material Impacts on Biofilm Formation

Crystal violet biofilm assays were used to analyze the biofilm-forming ability of *E. coli* and *B. subtilis* on the seven characterized materials, shown in Figure 2. *E. coli* and *B. subtilis* were selected for additional testing since they had the highest and lowest initial attachment, respectively (*E. coli* – 0.029 ± 0.0029; *B. subtilis* – 0.0056 ± 0.0023). Biofilm formation differed significantly between bacterial species and among materials (Table S.6.3).

**Figure 2:**
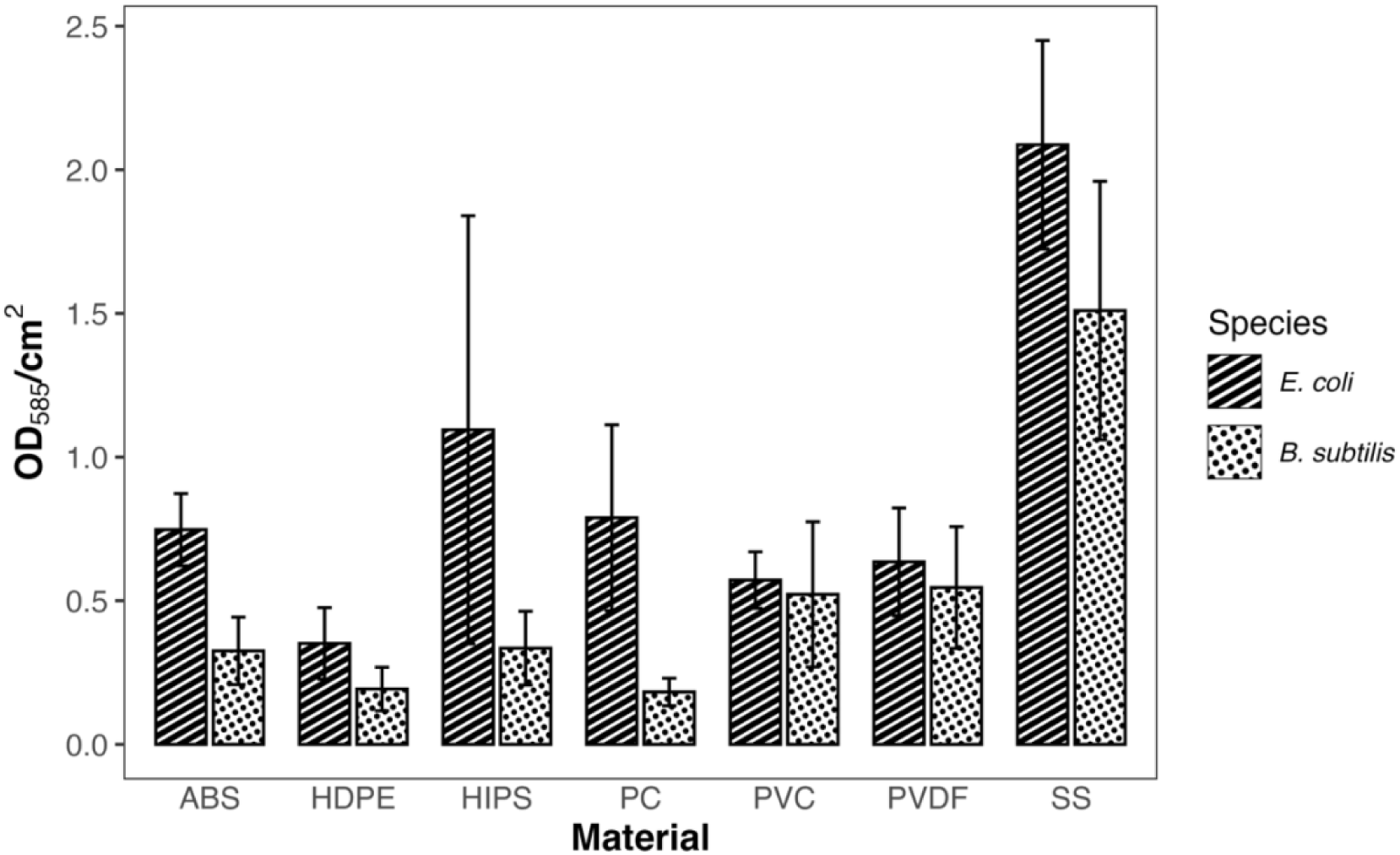
Surface area-normalized biofilm formation on different materials by *E. coli* and *B. subtilis*. Error bars represent the standard error from triplicate measurements.

Interestingly, material properties impacted biofilm formation, unlike initial attachment. Bonferroni-adjusted pairwise comparisons showed that stainless steel supported significantly greater biofilm biomass than the plastic materials (Table S.6.4). These results are particularly interesting, as we found that the highest biofilm formation occurred on the material with the lowest roughness and most hydrophilic surface. These results appear to run counter to trends reported previously in the literature^48,63^, where the greatest biofilm formation has been typically associated with rougher, more hydrophobic surfaces. Bohnic et al. found that biofilm formation, measured as crystal violet (CV) absorbance, increased with increasing surface roughness for surfaces with roughness ranging from 0.07µm to 5.8µm^48^. For *E. coli*, CV absorbance increased from 0.0267 ± 0.024 to 0.3117 ± 0.054. This trend was also seen for *P. aeruginosa* and *S. aureus*, with increases from 0.0517 ± 0.072 to 0.9417 ± 0.11, and 0.0537 ± 0.019 to 0.2697 ± 0.068, respectively^48^.

One explanation for this apparently contradictory result is the possible alteration of material surfaces by extracellular polymeric substances (EPS) produced during biofilm formation. Indeed, EPS constitutes a major component of biofilm biomass^64^ and contributes substantially to the biomass measured by the crystal violet assay^65,66^. We test this hypothesis using an adsorption experiment performed using dextran as a polymer representative of EPS (Figure S.7.1). Results correlate well with biofilm formation, with EPS adsorption showing a preferential affinity toward SS materials. A second hypothesis is that the SS caused stress to the bacterial cells when in contact with the stainless steel surface. Bacteria have been shown to exhibit increased biofilm formation in response to stressors^67,68^. EPS produced in the formation of a biofilm protects cells from stresses^69,70^. In our system, we hypothesize that the induction of a stress response when in contact with stainless steel resulted in elevated production of EPS.

*E. coli* biofilm formation was greater than that of *B. subtilis*, similar to results seen for initial attachment. The relative difference between *E. coli* and *B. subtilis* was greater for initial attachment than for biofilm formation, with *E. coli* exhibiting approximately 6.4-fold higher attachment and 1.7-fold higher biofilm formation. *B. subtilis* is known to be a strong biofilm former^71,72,73^. On the other hand, *E. coli* is less known for its biofilm-forming ability, although biofilm formation is strongly influenced by environmental and genetic factors^74^. Despite the fact that *B. subtilis* is typically a stronger biofilm former than *E. coli*, the significant difference in initial attachment seen in this study may drive lower biofilm formation on the studied materials.

## ENVIRONMENTAL IMPLICATIONS

Microbial biofilms serve as pathogen reservoirs in engineered water systems, including hospital plumbing. This work showed that bacterial properties, rather than material properties, drive the initial colonization of materials, and that the initial attachment of bacteria is relatively independent of subsequent biofilm development on the surfaces tested in this work. Longer-term biofilm colonization remained dependent on bacterial properties, with some material-specific dependence. This material dependence was associated with preferential adsorption of polysaccharides to particular material surfaces, which suggests that surface conditioning of surfaces in the environment may play a greater role in biofilm colonization. These results suggest that material selection, or modification, to limit initial bacterial attachment may not be the most effective strategy to reduce the presence of pathogenic biofilms. Rather, strategies which account for bacterial properties and longer-term surface conditioning may provide more effective approaches for managing pathogens in engineered systems. Taking advantage of bacterial properties to establish beneficial microbial communities may offer a promising alternative strategy for future investigation.

## ASSOCIATED CONTENT

### Supporting Information

Experimental details and methods, including a schematic of experimental setup (Figures S.1.1 – S.3.1; Tables S.2.1 – S.4.1); additional material characterizations (Figures S.5.1 – S.5.3); statistical results for two-way ANOVA and Bonferroni post-hoc test results (Tables S6.1 – S.6.14); adsorption of dextran (Figure S.7.1); relationship between material surface properties and bacterial attachment efficiency (Figure S.8.1).

### Funding Sources

This work was supported primarily by the Engineering Research Centers Program of the National Science Foundation under NSF Cooperative Agreement No. EEC-2133504. This work was performed in part at the Duke University Shared Materials Instrumentation Facility (SMIF), a member of the North Carolina Research Triangle Nanotechnology Network (RTNN), which is supported by the National Science Foundation (award number ECCS-2025064) as part of the National Nanotechnology Coordinated Infrastructure (NNCI).

### Notes

The authors declare no competing financial interest.

## ACKNOWLEDGEMENT

We thank Claudia Gunsch and her laboratory (Duke) for providing the *E. coli* and *B. subtilis* strains, and Deverick Anderson and his laboratory (Duke) for providing the *P. aeruginosa* and S*. aureus* strains.

