## Supporting Information for "The Influence of Material Surface Properties on Initial Bacterial Attachment and Biofilm Formation on Built Environment Surfaces"

Number of Pages: 14

Number of Figures: 7

Number of Tables: 7

S.1. TRACER EXPERIMENT INFORMATION

Tracer experiments were performed as described in the Methods section in the associated publication. A schematic of the column setup used in all three types of column experiments (bacteria, tracer, positive particles) is shown in Figure S.1.1. Tracer tests were carried out using 10mM KNO_3_, in which absorbance was measured at 304nm.

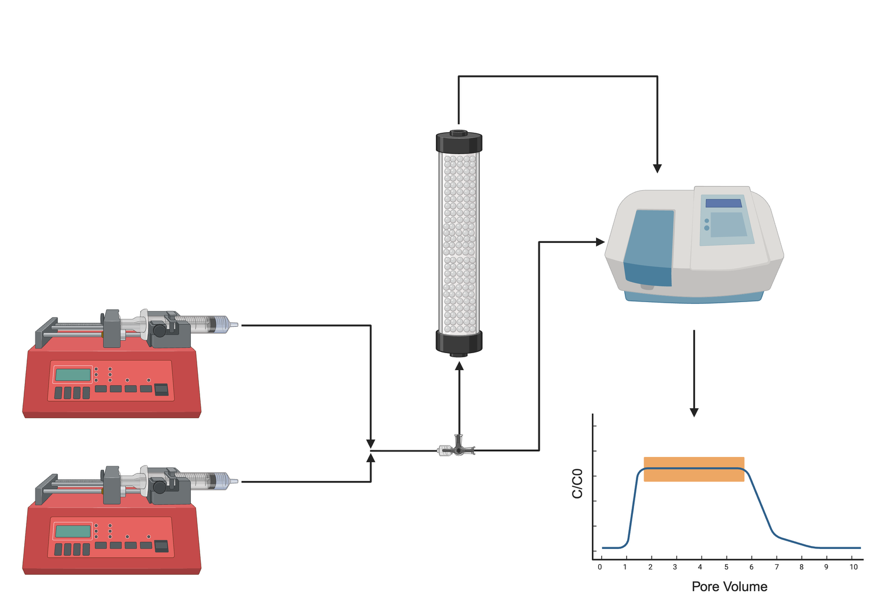

**Figure S.1.1:** Illustration of column experiment setup. Created in BioRender. Talma, K. (2025) <https://BioRender.com/quuyobp>

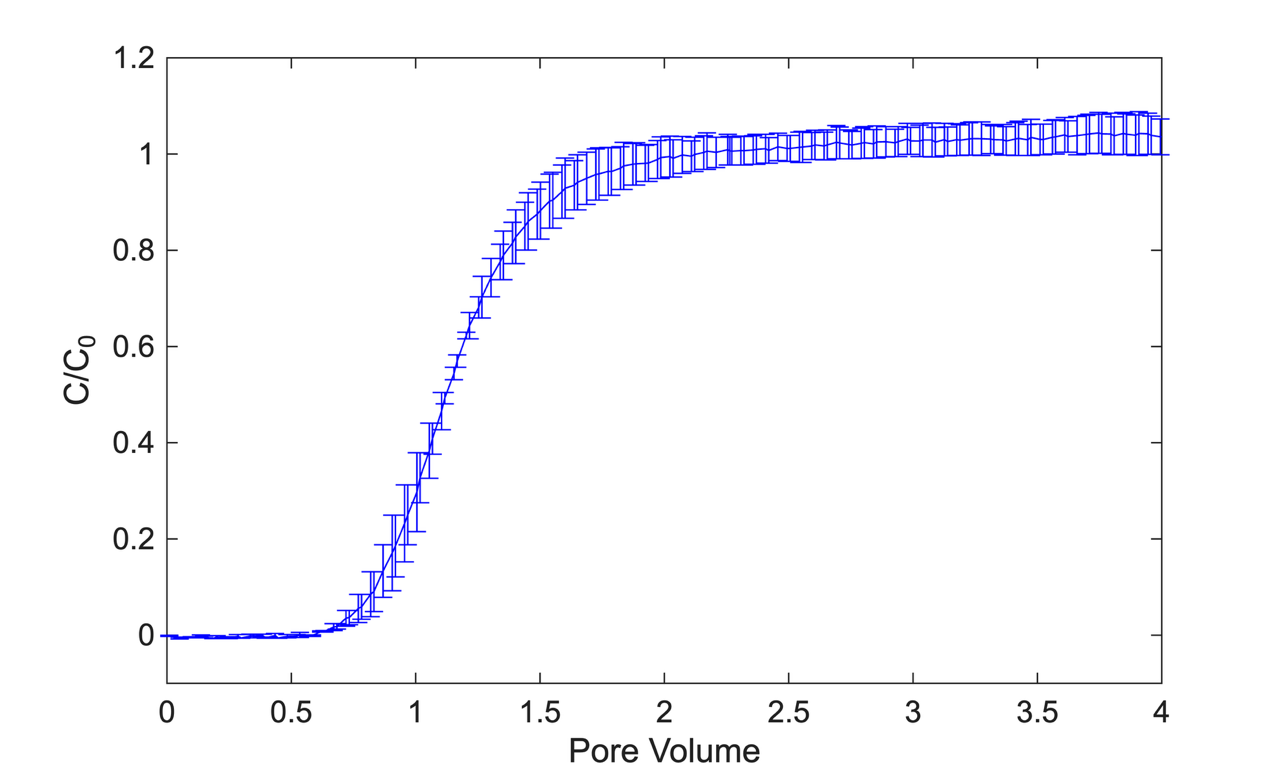

**Figure S.1.2:** Sample Tracer Test Plot. Potassium nitrate is used as the tracer at 10 mM and at a 2.95 mL/min superficial velocity. Error bars are standard error for two replicates. Plateau is achieved at 2 pore volumes.

S.2. POSITIVE PARTICLE COLUMNS AND CALCULATION OF ALPHA

Columns were carried out with aminated silica particles (nanoComposix; San Diego, California) to calculate the attachment efficiency (α) of bacteria using the normalization equation (Eq. 2). The positive particle experiments were performed as described in the Methods section with the following differences: (1) the column was packed to the 5cm line with material, (2) DI water was used as the background solution, and (3) absorbance was measured at 350nm. The C/C_0_ values reported in Table S.2.1 show that these particles have a high affinity to the test materials.

**Table S.2.1:** Measured Steady State Values from Breakthrough Curves

| **Material** | **C/C_0_*^a^*** |
| --- | --- |
| ABS | 0.00913 ± 0.000200 |
| HDPE | 0.0133 ± 0.00187 |
| HIPS | 0.00483 ± 0.000899 |
| PC | 0.0129 ± 0.00300 |
| PVC | 0.00584 ± 0.0000944 |
| PVDF | 0.0346 ± 0.0105 |
| SS | 0.0282 ± 0.00573 |

*^a^*Error is standard error based on duplicate measurements.

The predicted single-collector efficiency (η_0_) was calculated using the equation described by Tufenkji & Elimelech (2004)^39^:

$\eta_{0}=2.4A_{s}^{1/3}N_{R}^{-0.081}N_{Pe}^{-0.715}N_{vdW}^{0.052}+0.55A_{s}N_{R}^{1.675}N_{A}^{0.125}+0.22N_{R}^{-0.24}N_{G}^{1.11}N_{vdW}^{0.053}$ where the dimensionless parameters are defined in Table S.2.2.

**Table S.2.2:** Dimensionless Parameters used in Predicted Single-Collector Efficiency (η_0_) Calculation

| **Parameter** | **Definition^a^** | **Physical Interpretation** |
| --- | --- | --- |
| *A_s_* | $\frac{2\left( 1- \gamma^{5} \right)}{2-3\gamma+3\gamma^{5}-2\gamma^{6}}$ | Porosity-dependent parameter |
| *N_R_* | $\frac{d_{p}}{d_{c}}$ | Aspect ratio |
| *N_Pe_* | $\frac{Ud_{c}}{D}$ | Peclet number |
| *N_vdW_* | $\frac{A}{kT}$ | van der Waals number |
| *N_A_* | $\frac{A}{12\pi\mu a_{p}^{2}}$ | Attraction number |
| *N_G_* | $\frac{2}{9}\frac{a_{p}^{2}\left( \rho_{p}-\rho_{f} \right)g}{\mu U}$ | Gravity number |

^a^The parameters in the dimensionless parameters are: γ = (1 - ε)^1/3^, ε is the porosity, *d_p_* is the particle diameter [m], *d_c_* is the collector diameter [m], *U* is the fluid approach velocity [m s^-1^], *D* is the bulk diffusion coefficient as described by the Stokes-Eistein equation [m^2^/s], *A* is the Hamaker constant [J], *k* is the Boltzmann constant [J K^-1^], *T* is the fluid absolute temperature [K], *µ* is the absolute fluid viscosity [kg m^-1^ s^-1^], *a_p_* is the particle radius [m], *ρ_p_* is the particle density [kg m^-3^], *ρ_f_* is the fluid density [kg m^-3^], and *g* is the gravitational acceleration [m s^-2^].

S.3. BACTERIAL ATTACHMENT EXPERIMENTS – BREAKTHROUGH CURVES

Breakthrough curves, in which bacterial cells become observable in the column effluent and measured by UV-Visible spectrophotometer, are shown below.

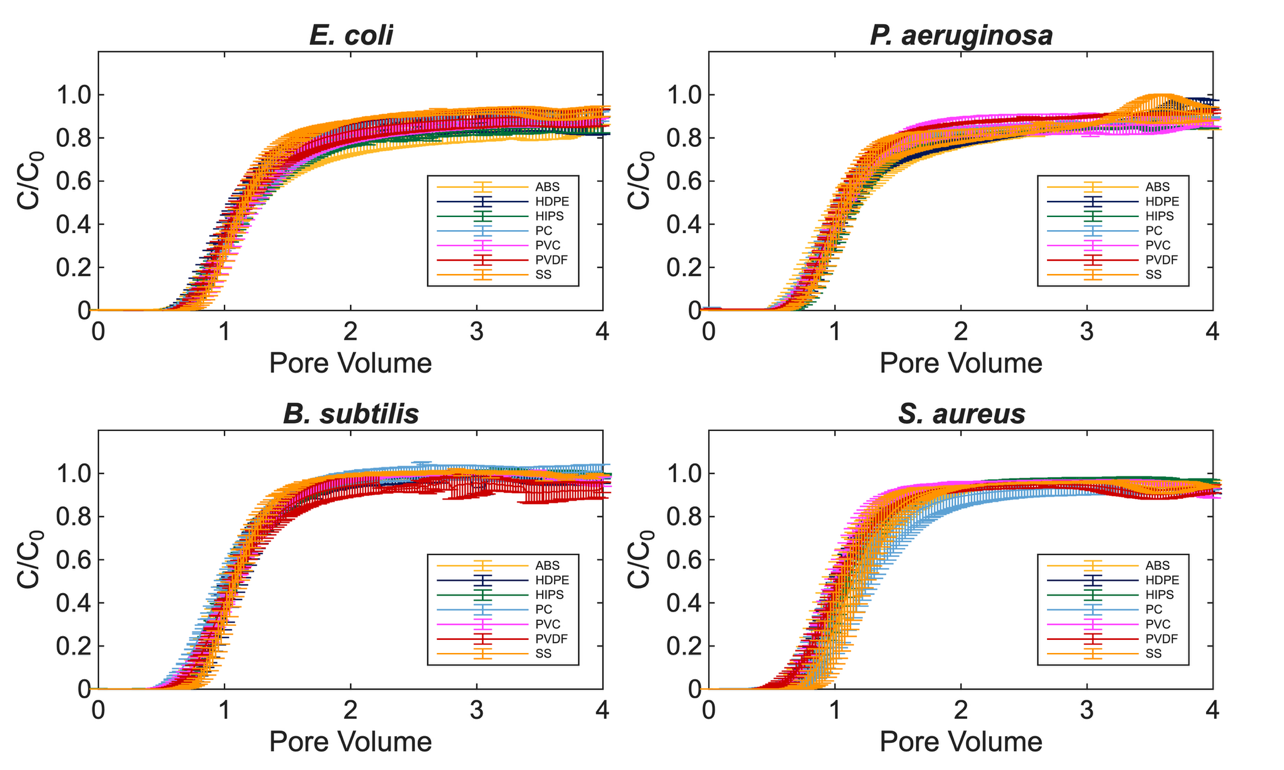

**Figure S.3.1:** Breakthrough curves for the 4 bacteria as a function of material. Breakthrough curves are measured at pH 7.5. The error bars are the standard error for duplicate measurements.

S.4. EPA MODERATELY HARD WATER PREPARATION

The following protocol was used to prepare 1 L of EPA Moderately Hard Water. 0.95 L of MILLI-Q, or equivalent, deionized water was placed in a properly cleaned carboy. 60 mg of MgSO_4_, 96mg NaHCO_3_, and 4mg KCl were added to the carboy and aerated overnight. 60mg CaSO_4_•2H_2_O was added to 0.05 L of MILLI-Q or equivalent deionized water in a separate flask. This flask was stirred on a magnetic stirrer until calcium sulfate was dissolved and added to the aerated 0.95 L and mixed well. The combined solution was aerated vigorously for an additional 24 hours to dissolve the added chemicals and stabilize the solution. Final prepared water quality parameters are shown in Table S.4.1.

**Table S.4.1:** Preparation of Synthetic Freshwater using Reagent Grade Chemicals^1^

|  | **Reagent Added (mg/L)^2^** | | | | **Approximate Final Water Quality** | | |
| --- | --- | --- | --- | --- | --- | --- | --- |
|  | NaHCO_3_ | CaSO_4_×2H_2_O | MgSO_4_ | KCl | pH^3^ | Hardness^4^ | Alkalinity^4^ |
| Moderately Hard | 96 | 60 | 60 | 4 | 7.4 -7.8 | 80 - 100 | 57 - 64 |

^1^Taken in part from Marking and Dawson (1973). ^2^Add reagent-grade chemicals to deionized water. ^3^Approximate equilibrium pH after 24 hours of aeration. ^4^Expressed as mg CaCO_3_/L.

S.5. ADDITIONAL CHARACTERIZATION OF SURFACES

Images obtained from 3D optical/ laser confocal profilometry were used to determine the surface roughness of the materials tested.

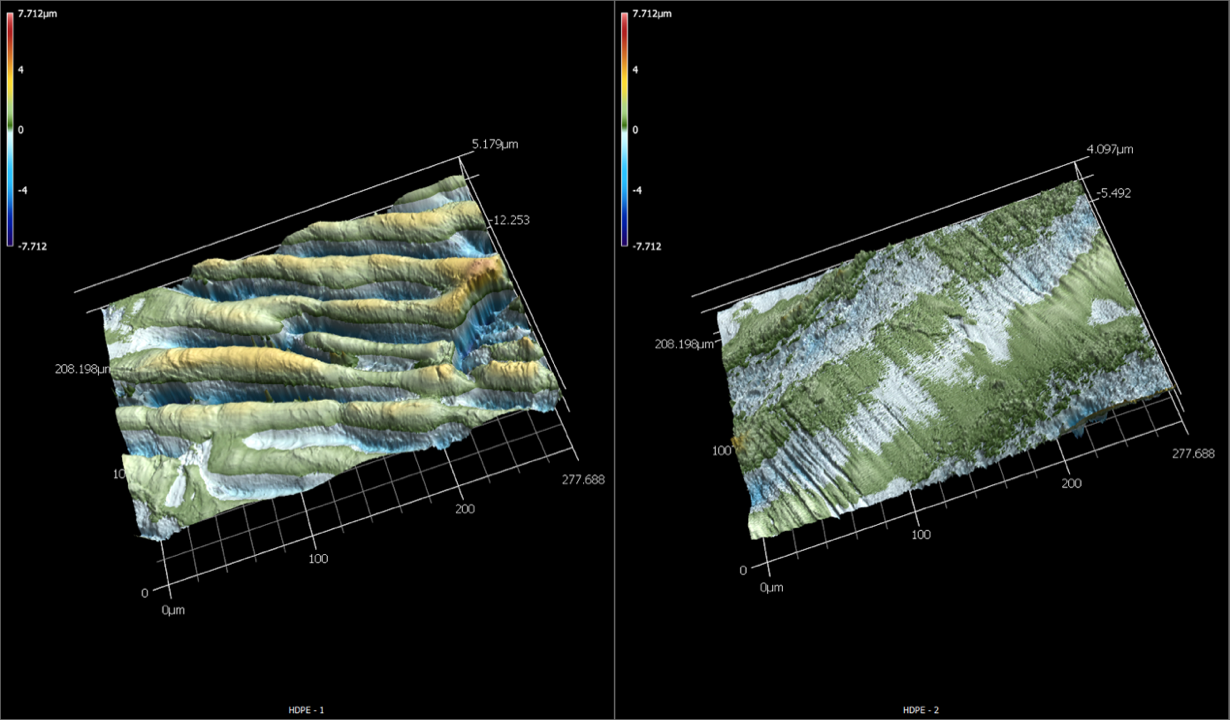

**Figure S.5.1:** Three-dimensional display of 3D optical/ laser confocal profilometry measurements taken on the surface of two HDPE samples, illustrating the heterogeneity of the surfaces.

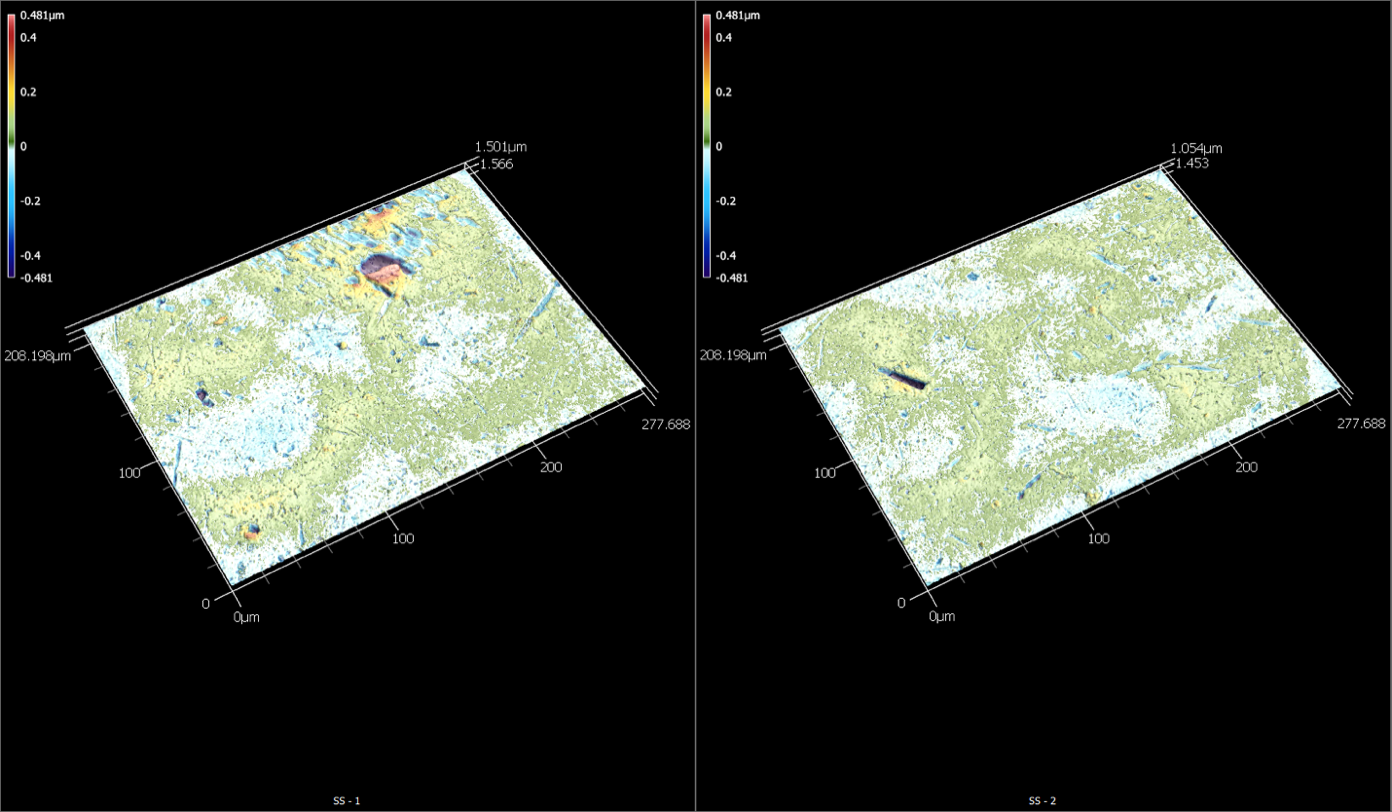

**Figure S.5.2:** Three-dimensional display of 3D optical/ laser confocal profilometry measurements taken on the surface of two SS samples, illustrating the heterogeneity of the surfaces.

The FTIR spectra collected are shown below.

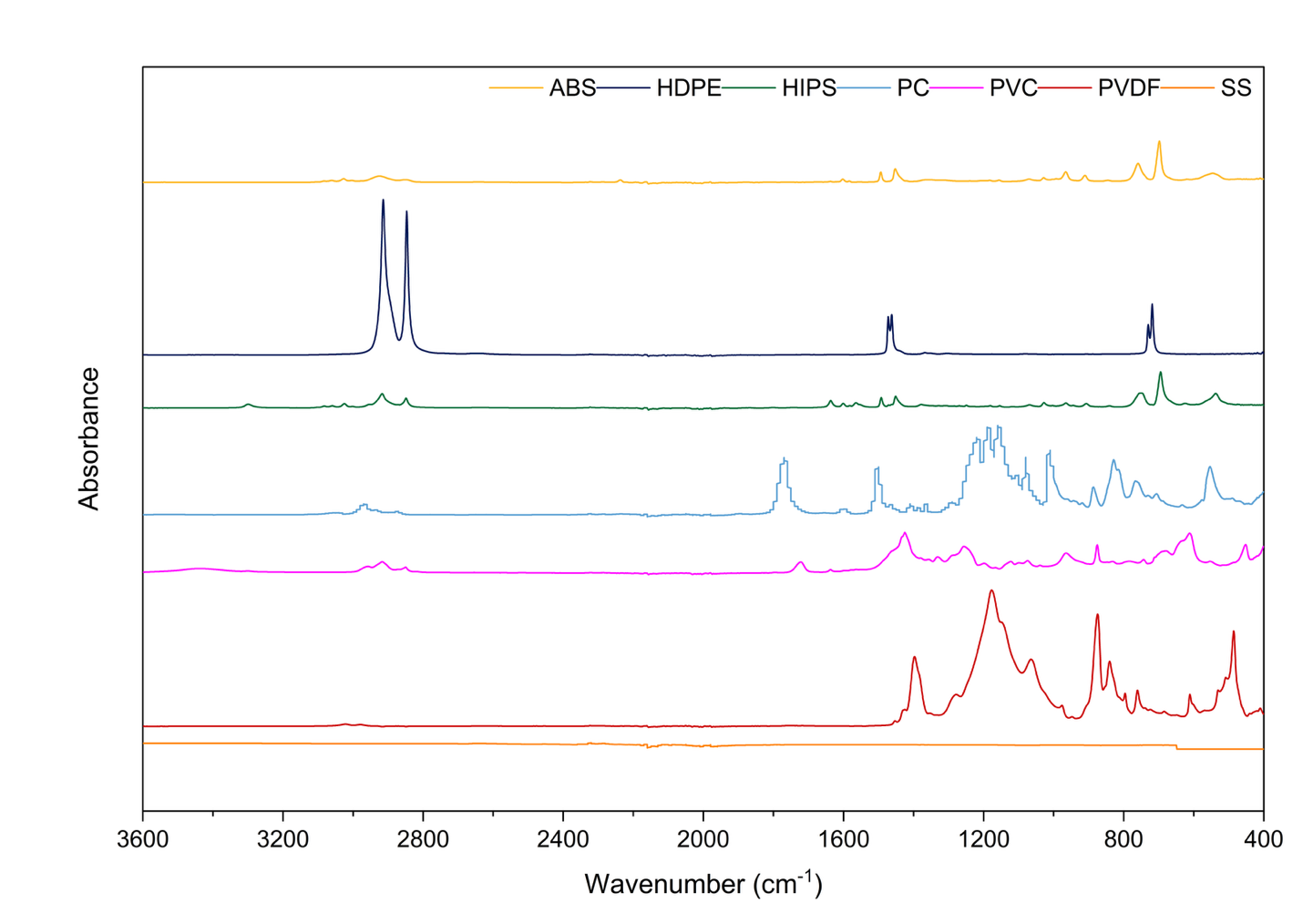

**Figure S.5.3:** FTIR spectra of ABS, HDPE, HIPS, PC, PVC, PVDF, and SS in the spectral range 3600 – 400 cm^-1^.
S.6. RESULTS OF STATISTICAL TESTS

**Table S.6.1:** Two-way ANOVA evaluating the effects of material, bacterial species, and their interaction on attachment efficiency (α).

| **Source** | **df** | **Sum of Squares** | **Mean Square** | **F** | **p-value** |
| --- | --- | --- | --- | --- | --- |
| Material | 6 | 0.000848 | 0.000141 | 2.2331 | 0.069291 |
| Species | 3 | 0.005446 | 0.001815 | 28.67685 | 1.1 x 10^-8^ |
| Material x Species | 18 | 0.000567 | 3.15 x 10^-5^ | 0.497321 | 0.937275 |

**Table S.6.2:** Bonferroni-adjusted pairwise comparisons of estimated marginal means among bacterial species following a significant main effect of species.

| **Group 1** | **Group 2** | **Difference** | **Adjusted p-value** | **Confidence Interval** |
| --- | --- | --- | --- | --- |
| *E. coli* | *P. aeruginosa* | 0.002422 | 1 | [-0.006, 0.011] |
| *E. coli* | *B. subtilis* | 0.023005 | 1.48 x 10^-7^ | [0.014, 0.032] |
| *E. coli* | *S. aureus* | 0.018094 | 1.05 x 10^-5^ | [0.010, 0.027] |
| *P. aeruginosa* | *B. subtilis* | 0.020584 | 1.17 x 10^-6^ | [0.012, 0.029] |
| *P. aeruginosa* | *S. aureus* | 0.015672 | 9.33 x 10^-5^ | [0.007, 0.024] |
| *B. subtilis* | *S. aureus* | -0.00491 | 0.681589 | [-0.013, 0.004] |

**Table S.6.3**: Two-way ANOVA evaluating the effects of material, bacterial species, and their interaction on biofilm biomass.

| **Source** | **df** | **Sum of Squares** | **Mean Square** | **F** | **p-value** |
| --- | --- | --- | --- | --- | --- |
| Material | 6 | 8.723664 | 1.453944 | 5.39352 | 0.000901 |
| Species | 1 | 1.527417 | 1.527417 | 5.666073 | 0.024617 |
| Material x Species | 6 | 0.702598 | 0.1171 | 0.43439 | 0.84936 |

**Table S.6.4:** Bonferroni-adjusted pairwise comparisons of estimated marginal means among materials following a significant main effect of material.

| **Group 1** | **Group 2** | **Difference** | **Adjusted p-value** | **Confidence Interval** |
| --- | --- | --- | --- | --- |
| ABS | HDPE | 0.264314 | 1 | [-0.802, 1.330] |
| ABS | HIPS | -0.17849 | 1 | [-1.183, 0.826] |
| ABS | PC | 0.050814 | 1 | [-0.954, 1.056] |
| ABS | PVC | -0.01059 | 1 | [-1.016, 0.994] |
| ABS | PVDF | -0.0543 | 1 | [-1.059, 0.951] |
| ABS | SS | -1.26206 | 0.005318 | [-2.267, -0.257] |
| HDPE | HIPS | -0.4428 | 1 | [-1.509, 0.623] |
| HDPE | PC | -0.2135 | 1 | [-1.279, 0.852] |
| HDPE | PVC | -0.27491 | 1 | [-1.341, 0.791] |
| HDPE | PVDF | -0.31861 | 1 | [-1.385, 0.747] |
| HDPE | SS | -1.52637 | 0.001093 | [-2.592, -0.460] |
| HIPS | PC | 0.229303 | 1 | [-0.776, 1.234] |
| HIPS | PVC | 0.167896 | 1 | [-0.837, 1.173] |
| HIPS | PVDF | 0.124189 | 1 | [-0.881, 1.129] |
| HIPS | SS | -1.08357 | 0.025507 | [-2.089, -0.079] |
| PC | PVC | -0.06141 | 1 | [-1.066, 0.944] |
| PC | PVDF | -0.10511 | 1 | [-1.110, 0.900] |
| PC | SS | -1.31287 | 0.003382 | [-2.318, -0.308] |
| PVC | PVDF | -0.04371 | 1 | [-1.049, 0.961] |
| PVC | SS | -1.25147 | 0.005843 | [-2.256, -0.246] |
| PVDF | SS | -1.20776 | 0.008606 | [-2.213, -0.203] |

S.7. ADSORPTION OF DEXTRAN TO MATERIALS

Adsorption experiments were performed with dextran (Dextran from *Leuconostoc* spp., Sigma Life Science; St. Louis, Missouri) to assess the ability of extracellular polymeric substances (EPS) to adsorb to the characterized materials. To conduct the experiments, a known mass of the test material was added to 10mL of 15mg/L dextran solution and incubated at 25 °C for 24 hours. After incubation, the residual dextran solution was removed from each vial and measured using a total organic carbon (TOC) analyzer (Shimadzu; Columbia, Maryland). The adsorbed polymer concentration Γ (mg/cm^2^) was calculated according to Equation S.6.1:

$\Gamma= \frac{\left( C_{0}- C_{e} \right)\times V}{A}$	Eq. S.6.1

where C_0_ (mg/L) is the initial dissolved polymer concentration, C_e_ (mg/L) is the final dissolved concentration in tubes containing material, V (L) is the solution volume, and A (cm^2^) is the surface area of the material.

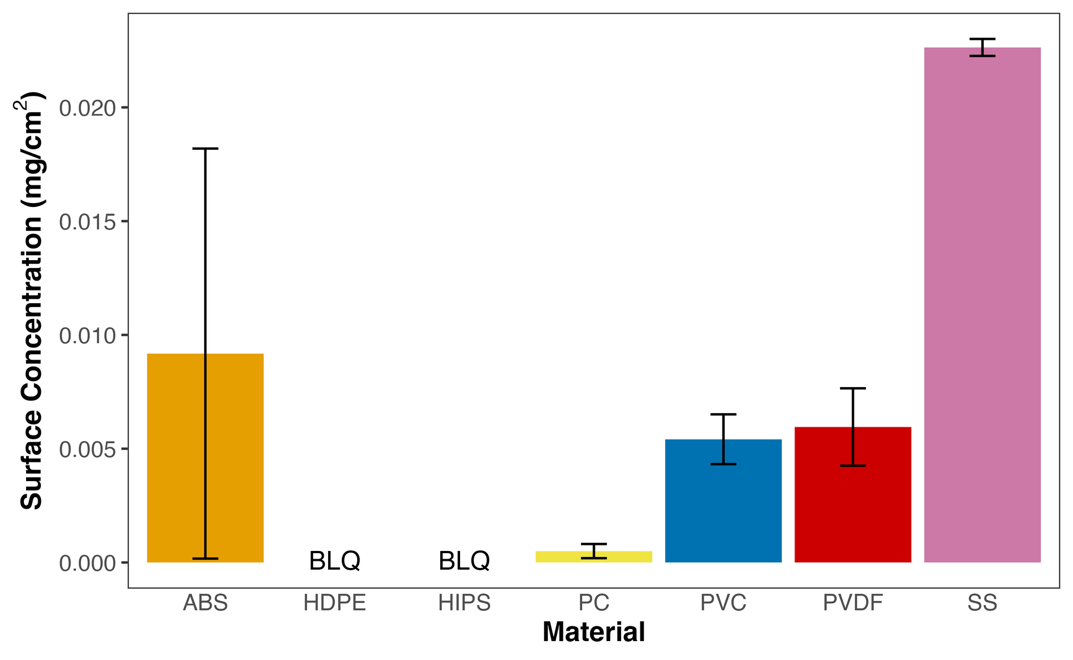

**Figure S.7.1:** Adsorption of dextran to material surfaces. Error bars represent the standard error from triplicate measurements. BLQ – below the limit of quantification.

S.8 RELATIONSHIP BETWEEN MATERIAL SURFACE PROPERTIES AND INITIAL ATTACHMENT EFFICIENCY (α)

Scatter plots were used to visualize the dependence of initial attachment efficiency (α) on the measured material properties.

| A | B |
| --- | --- |
| 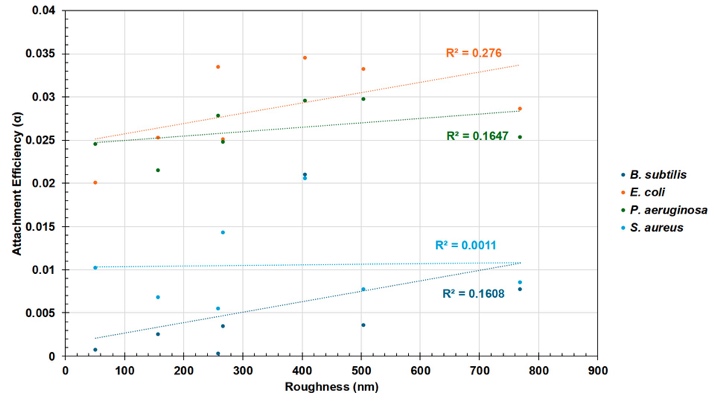 | 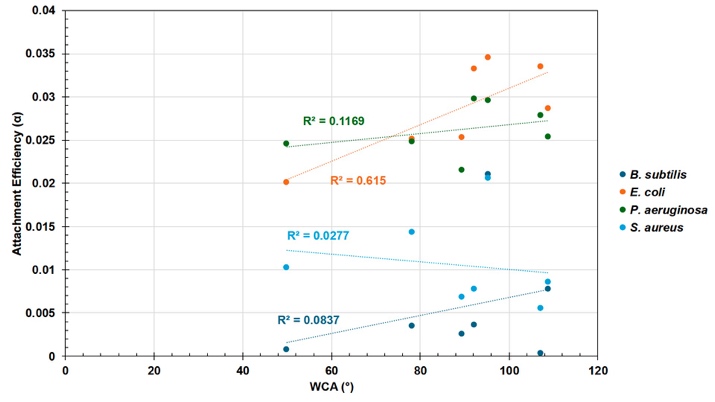 |
| C |  |
| 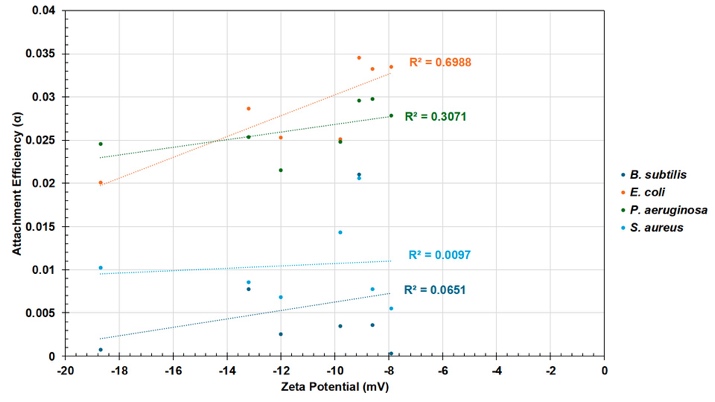 | |

**Figure S.8.1:** A) **Relationship between roughness and attachment efficiency for bacterial species. B) Relationship between WCA and attachment efficiency for bacterial species. C) Relationship between zeta potential (mV) and attachment efficiency. Dashed lines represent linear regressions for each bacterial species, with corresponding R^2^ values shown.**
